# Solid-state fermentation of oyster mushroom by-products using *Neurospora crassa*: a sustainable approach for the development of novel meat analogues

**DOI:** 10.64898/2026.04.30.721925

**Authors:** Pablo Navarro-Simarro, Bryan Moreno-Chamba, Julio Salazar-Bermeo, Lourdes Gómez-Gómez, Ángela Rubio-Moraga, Alberto López-Jimenez, Nuria Martí, Oussama Ahrazem

**Author notes:** Both authors equally contributed to this work (Pablo Navarro-Simarro, Bryan Moreno-Chamba). **Competing interests** The authors declare no competing interest. **Corresponding Author** Oussama Ahrazem.

## Abstract

Mushroom production generates large amounts of by-products, particularly stipes, which can represent up to half of the fruiting body biomass. Due to their similar composition to mushroom caps, these residues represent a promising substrate for the development of value-added foods. In this study, oyster mushroom stipes were used as a substrate for solid-state fermentation (SSF) with a *Neurospora crassa* strain isolated in Albacete to produce a novel meat analogue inspired by the *oncom*. Fermentation generated a cohesive matrix bound by hyphae that adopted the shape of the mold and exhibited a meat-like color, although with a softer texture. Nutritional analysis revealed a product with relatively low protein content but a complete amino acid profile, enriched in dietary fiber and containing unsaturated fatty acids. These results demonstrate that SSF with *N. crassa* provides a strategy to upcycle oyster mushroom by-products into fiber-rich meat analogues with potential applications in sustainable food systems.

## 1. Introduction

In recent years, solid state fermentation (SSF) has gained a lot of interest for the upcycling of agricultural by-products into microbial protein, enzymes, pigments or novel foods (Fernandez Castaneda et al., 2025; Makran et al., 2026; Yafetto, 2022). This process is of lower cost than submerged fermentation because it requires less water, fewer equipment and there is a minor risk of microbial contamination (Wang et al., 2023). Among the micro-organisms used for SSF, the use of fungi of the genus *Neurospora* stands out due to the interest of researchers in using indigenous edible fungal strains. Specifically, *Neurospora intermedia* is valued in Indonesia for its ability to convert tofu by-products or bagasse from peanut oil extraction into red *oncom*, a traditional fermented food. Recently, some studies have used *N. intermedia* to perform SSF on agro-industrial plant residues such as bagasse, hulls or seeds to obtain novel foods similar to *oncom* (Gmoser et al., 2020; Maini Rekdal et al., 2024).

A similar fungus, *Neurospora crassa*, inhabits burnt wood or other lignocellulosic wastes (Kuo et al., 2014), although it was first described in 1843 as a contamination agent in French bakeries (Roche et al., 2014). This fungus has potential for degrading agro-industrial waste because it has genes for degrading polysaccharides such as cellulose, hemicellulose or mannan (Samal et al., 2017). In the field of nutrition, *Neurospora* has been shown to be a fungus suitable for human consumption and has great potential for formulating alternatives to meat (Bartholomai et al., 2022). Its safety lies in the fact that it does not express mycotoxins and there are no reports of human allergens (Lozy et al., 2022). Moreover, *N. crassa* strains are a source of protein and prebiotic fibre (Gmoser et al., 2019; Huang et al., 2022), increase the digestibility of the substrate (Li et al., 2019) and enrich the final product with higher levels of vitamin E and vitamin D2 (Gmoser et al., 2020).

Among the agricultural by-products that could be revalued by *Neurospora*, mushroom stem waste is among the least studied even though it accounts to 30% of fruiting body biomass (Guo et al., 2022). Moreover, the global mushroom market was estimated at 17.25 million tonnes in 2023, and it is expected to reach 32.04 million tonnes by 2032 (*Mushroom Market Size, Share, Growth, Analysis, 2032*, n.d.), which will involve the management of a large amount of by-products. Nutritionally, the composition of the stipes is very similar to that of the commercial mushroom cap, rich in polysaccharides such as β-glucans and chitin, some of which have a prebiotic effect (Navarro-Simarro et al., 2024, 2025; Törős et al., 2023). *Pleurotus ostreatus*, one of the most commercialized mushrooms, is also interesting because it has a significant content of essential amino acids, as well as a good protein digestibility (Carrasco-González et al., 2017). The combination of the nutritional value of edible mushrooms combined with *N. crassa* fermentation may promote a new range of nutritious meat alternatives aligned with the Circular Economy.

Given the above, this study aims to revalorize oyster mushroom (*P. ostreatus*) stems with a strain of *N. crassa* isolated in Albacete (Spain) for the formulation of vegan patties. The effect of SSF on the structure, nutritional composition, functional and sensory aspects were assessed. The formulation of new foods based on *Neurospora* is not only a way to obtain new tasty or functional products, but also to value the great variety of traditional fermented foods that exist in Indonesia promoting circular economy in mushroom production.

## 2. Materials and Methods

### 2.1 Chemicals and reagents

Potato Dextrose Broth (PDB) and glycerol (≥99%) were obtained from Scharlab (Barcelona, Spain), while bacteriological agar (European grade) was purchased from Conda (Madrid, Spain). Corn oil was sourced from Deoleo Global S.A. (Alcolea, Spain). A commercial pork and beef burger, produced by Centros Comerciales Carrefour S.A. (Alcobendas, Spain), was included for comparison. Additionally, tempeh was supplied by Master Terra Verda S.L. (Valencia, Spain). Reagents for *in vitro* digestion as well as bacterial culture media for bacterial growth stimulation experiments were purchased from Merck (Madrid, Spain) and Equilabo Scientific S.L. (Murcia, Spain), respectively.

### 2.2 Mushroom by-products

The oyster mushroom stipes were supplied by the company Setas Meli S.L. derived from the cultivation of edible mushrooms in Casasimarro (Spain). The by-products were received within 24 h after harvesting, washed, and stored at -20°C. Afterwards, they were freeze-dried at -50°C for 72 h (LyoQuest-85 / 208 V 60 Hz, Teslar). Then, the samples were milled, and the powder was rehydrated to 300% of its weight with distilled water. The freeze-dried substrate, used to simulate the starting point as fresh product, was finally rehydrated, autoclaved, and stored at 4°C until inoculation. The oyster mushroom powder is referred in the text as OM, while the autoclaved version is labeled as OM-sterilized.

### 2.3 *Neurospora crassa* strain

The *Neurospora crassa* strain utilized in this study was originally isolated from decaying plant debris collected in Albacete, Spain. The strain was cultured on potato dextrose agar (PDA) and taxonomically identified through the amplification of its internal transcribed spacer (ITS) regions, followed by sequence comparison using the BLASTn algorithm (Navarro Simarro & López-Jiménez, 2026). Spore suspensions of the fungus were prepared in 20% (v/v) glycerol and stored at -80°C for long-term preservation. Prior to experimental use, the strain was reactivated by cultivation on PDA or directly placed in sterilized mushroom substrate. This strain was registered in the Spanish Type Culture Collection as *Neurospora crassa* CECT 21312.

### 2.4 Solid-state fermentation and product formulation

Briefly, 60 g of sterilized and hydrated mushroom substrate was placed in containers measuring 10 cm in diameter and 14 cm in height, each covered with a mesh lid to allow oxygen exchange. The substrate was then inoculated with 2 mL of a suspension containing *N. crassa* mycelium and spores resuspended in distilled water (10^8^ propagules/mL). Fermentation was carried out for 48 hours at 28°C. Following fermentation, the raw product was removed from the container and refrigerated at 4°C until further use. Subsequent cooking involved a stir-fry process using olive oil at 180°C for 5 minutes per side. The fermented oyster mushroom with *N. crassa* was referred to as Ferm-OM, while the stir-fried version was labeled as Ferm-OM-cooked.

As a proof of concept for pilot scale-up, fermentation was also carried out in polyethylene bags with holes to allow gas exchange and produce larger quantities of product. For this purpose, 100 g of moistened substrate were inoculated with *N. crassa* and the same parameters of incubation as above were used. This type of fermentation is typical of fungal fermented products such as tempeh (Santhirasegaram et al., 2016).

### 2.5 Color analysis of samples

Color measurements were performed at room temperature using a Minolta Chroma Meter CR-300 (Minolta Co., Ltd., Chuo-Ku, Osaka, Japan), a tristimulus colorimeter based on the CIE (Commission Internationale de l’Éclairage) L, a, b color system. The device was calibrated prior to use with a white reference tile (L = 96.94, a = +0.18, b = +1.89) (González et al., 2015). A block of commercial tempeh and a commercial beef patty (60% pork and 40% beef) were used for comparative.

From the measured L*, a*, and b* values, additional color parameters were calculated. Hue°, representing the color tone angle in the CIELAB plane, was calculated as (Eq. 1):

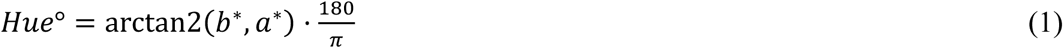

Chroma (C*), representing color saturation or intensity, was calculated as (Eq. 2):

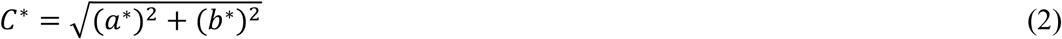

The color difference of each sample (ΔE) relative to the Meat cooked control was expressed as (Eq. 3):

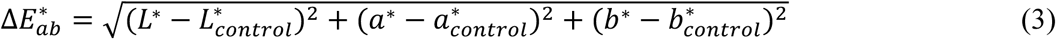

### 2.6 Field emission scanning electron microscopy (FESEM) of samples

Samples were frozen at -80°C and subsequently freeze-dried at -50°C for 48 h (LyoQuest-85 / 208 V 60 Hz, Teslar). Micrographs of the samples were captured by Sigma 300 VP FESEM (Carl Zeiss, Germany) at 20 kV without coating.

### 2.7 Technofunctional properties

Briefly, 0.33 g of freeze-dried sample was added to 10 mL of distilled water in a centrifuge tube and incubated at room temperature for 18 hours. To assess water absorption, the sample was centrifuged at 1250×*g* for 30 minutes, and for water retention, it was centrifuged at 3000×*g* for 20 minutes. The water absorption or water retention were determined by weighing the wet pellet and using the following formula (Eq. 4):

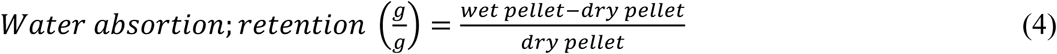

The supernatants obtained from the centrifugation step for water retention analysis were incubated and dried at 60°C for 24 hours in 5 cm diameter glass petri dishes. The precipitated solutes were then weighed to calculate solubility.

The oil retention capacity was tested using the protocol of Wang *et al*. (2020), with modifications (Wang et al., 2020). Briefly, 0.3 g of freeze-dried material was placed on a 50 mL tube followed by the addition of 15 mL of commercial corn oil. The mixture was incubated at room temperature for 20 minutes, mixing the suspensions for 15 seconds every 5 minutes. Afterward, the tubes were centrifugated at 1600×*g* for 25 minutes and the pellet is weighed. The oil retention capacity is determined by the following formula (Eq. 5):

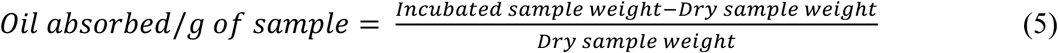

### 2.8 Texture analysis of samples

Textural properties of samples were evaluated using a texturimeter (TA.XTPlusC, Aname Instrumentación Científica, Madrid, Spain). Ferm-OM, Ferm-OM-cooked, raw tempeh, stir-fried tempeh and fermented mushroom in their raw and cooked versions were tested. Penetration was tested with a 20 mm (P/20) aluminum cylindrical probe to a depth of 10 mm, while compression was tested with a 75 mm (P/75) compression platform to a deformation of 5 mm. Parameters such as maximum force, positive and negative peak force and distance, positive and negative area under the force-time curve, slope at positive peak and force at 5 mm deformation were recorded. The data obtained were processed to calculate the mean, standard deviation and coefficient of variation of each experimental condition, allowing comparison of the firmness, strength and mechanical behavior of the different samples.

### 2.9 Nutrient analysis of samples

The compositional analysis was conducted by AINIA Centro Tecnológico (Paterna, Spain) using validated analytical protocols. Total amino acids were quantified by HPLC following lipid extraction, with cysteic acid used for cysteine quantification and methionine reported as methionine sulfone. Total fat was determined through gravimetric analysis after acid hydrolysis, while protein content was measured using the Dumas combustion method. Total dietary fiber was assessed via an enzymatic-gravimetric method. Total sugars and insoluble carbohydrates were analyzed volumetrically, with results expressed as glucose equivalents. The determination of insoluble carbohydrates includes starch and/or other insoluble polysaccharides, and results are expressed as a percentage of glucose. Salt content was determined using Inductively Coupled Plasma–Atomic Emission Spectroscopy (ICP-AES) and calculated as sodium × 2.5, following EU Regulation No. 1169/2011. Finally, the fatty acid profile was analyzed using Gas Chromatography with Flame Ionization Detection (GC-FID).

The calculation of the energy value (kcal) was based on Regulation (EU) No 1169/2011. This regulation assumes that carbohydrates and proteins provide 4 kcal/g, fiber provides 2 kcal/g, and fats provide 9 kcal/g. The formula used to calculate the energy value is (Eq. 6):

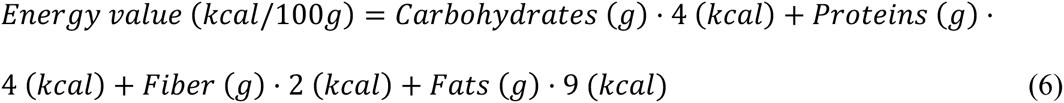

To calculate the moisture content, the fresh (S_wet_) and freeze-dried (S_dry_) raw samples were weighed and the following formula was used (Eq. 7):

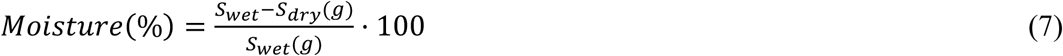

### 2.10 Prebiotic potential of samples

#### 2.10.1 *In vitro* digestion of samples

Prior the experiment, the samples were subjected to an *in vitro* digestion to assess their potential to stimulate bacterial proliferation selectively after ingestion, following the protocol established by Minekus et al. (Minekus et al., 2014). After subjecting the samples to oral, gastric and intestinal digestion phases, the digested samples were snap frozen and stored at -80°C until analysis. An undigested aliquot of each sample was also used in the experiments.

#### 2.10.2 Growth curve monitorization

Representative strains were tested for their ability to utilise mushroom polysaccharides as growth substrates using a microtiter plate method (Chung et al., 2017; Moreno-Chamba et al., 2025), with modifications. The bacterial strains used were purchased from the Spanish Type Culture Collection (CECT) such as: *Bifidobacterium bifidum* CECT 870, *B. longum* CECT 4503*, Lactobacillus brevis* CECT 4021, *Lactiplantibacillus plantarum* 748, *Lactococcus lactis* subsp. *lactis* CECT 185, *Pediococcus acidilactici* CECT 5911, and *Streptococcus salivarius* subsp. *thermophilus* CECT 7207. *Escherichia coli* CECT 515 was used as a pathogenic control.

The samples (OM, Ferm-OM, or Ferm-OM-cooked, both digested and undigested) were mixed separately with buffered peptone water in a 0.4% w/v, and then sterilized by autoclaving. Substrate controls of buffered peptone water supplemented with 0.4% w/v of glucose or fructooligosaccharides (FOS) were included. For the assays, fresh bacterial suspensions (optical density at 650 nm = 0.5 McFarland or 10^8^ CFU/mL) were prepared in buffered peptone water. Then, the bacterial suspensions were mixed 1:1 (v/v) with either samples or substrate controls (final volume of 200 µL, 0.2% of substrate). Bacterial suspensions in buffered peptone water with no substrate (negative control) and pure buffered peptone water (blank) were included as well. The optical density (650 nm) was measured in a microplate reader (Cytation™ 3 Cell Imaging Multi-Mode reader, BioTek Instruments, Inc, Winooski, VT, USA) each 10 min for 24 h at 37°C under aerobic or anaerobic conditions, depending of each strain requirement. After incubation, the plates were stored at -20°C for further analysis.

#### 2.10.3 Growth curve parameters

The growth after incubation and maximum growth were recorded from the growth curves. The prebiotic activity score (PAS) was also calculated according to Moreno-Chamba et al. (Moreno-Chamba et al., 2022), for each of the undigested and digested samples. The growth rate was determined according to Pirt (Pirt, 1975), in the strains where positive PAS were identified.

### 2.11 Microbiological analysis

Microbiological analyses were performed by Vitab Laboratorios (La Gineta, Spain) following standardized protocols. Aerobic mesophilic bacteria were assessed at 31°C according to UNE-EN ISO 4833-1. Total coliforms and *Escherichia coli* were enumerated using RAPID’E.coli 2 Agar. Coagulase-positive *Staphylococcus* was analyzed following UNE-EN ISO 6888-1. *Salmonella* spp. detection was conducted using UNE-EN ISO 6579-1:2017. *Listeria monocytogenes* was quantified using RAPID’L.mono Agar.

### 2.12 Statistical analysis

All samples were analyzed in triplicate, and the results are expressed as mean ± standard deviation. Statistical analyses were performed using GraphPad Prism 8.0.2 and R. Data were evaluated for normality and homogeneity of variances before applying appropriate statistical tests. For comparisons between two groups, differences were assessed using a two-sample t-test. When more than two groups were compared, one-way analysis of variance (ANOVA) followed by Tukey’s or Dunnett’s post hoc tests was applied. Significance was considered at *p < 0.05*.

## 3. Results

### 3.1 Macroscopic appearance

After fermentation, the Ferm-OM showed a compact appearance with two clearly differentiated parts, the upper and the lower (**Figure 1**). As for the lower part of the raw product, it can be observed that it adopts the shape of the mold due to the formation of *N. crassa* hyphae that maintain the structure. The upper part, in direct contact with oxygen, presents a velvety appearance and the orange color characteristic of carotenoid expression by *N. crassa*. Shorter fermentation times decrease the amount of spores and prevent the formation of orange spots, although the structure becomes less compact. The choice of substrate and filamentous fungus for SSF was based on a pilot test conducted prior to this study (**Supplementary Figure 1**).

**Figure 1:**
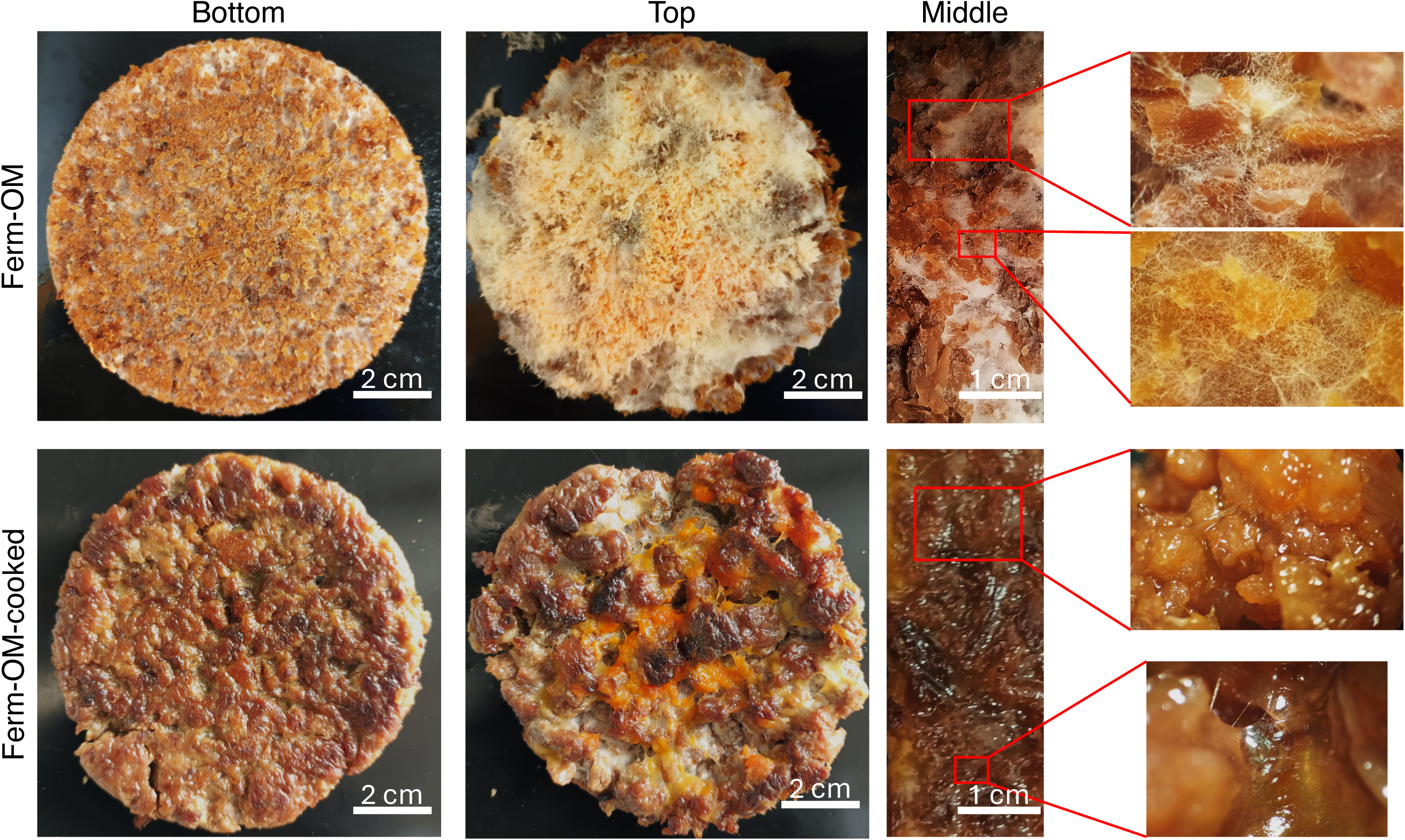
Macroscopic appearance of solid-state fermented oyster mushroom stem with *Neurospora crassa* after 48 h incubation at 28°C. The differences between the uncooked product (Ferm-OM) and the cooked product (Ferm-OM-cooked) can be observed. Images of the lower part of the product (bottom), in contact with the container, the upper part (top), where the aeration takes place, and the middle part are presented.

As for Ferm-OM-cooked, a noticeable color change of the mushroom hyphae to a dark brown is observed (**Figure 1**). The lower part adopts a palette of browns similar to that of the cooked meat, while the upper part shows some orange areas due to the cooking of the spore-rich zones. As for the longitudinal cuts, in the raw patty the granules coated and connected by the hyphae of *N. crassa* can be observed. Once cooked, these hyphae present a more viscous aspect, although they still connect the granules, and the overall color is brown as on the external surface. The preservation of the hyphae after cooking is a key aspect as it allows the patty to maintain its structure.

To evaluate the industrial feasibility of this product, batch fermentations of 100 g were carried out in perforated polyethylene bags (**Figure 2**). The external appearance of the product closely resembles that of *oncom*, featuring orange spots produced by *N. crassa* spores, which are primarily located around the aeration holes of the bag. Internally, the product is brown in color, with visible whitish hyphae of *N. crassa*. Notably, once the product comes into contact with oil, the white coloration disappears, and a cohesive brown block forms. After cooking, the spores appear as small dots corresponding to the aeration holes, which detach easily, facilitating cleaning. This proof of concept demonstrates that large-scale fermentations of oyster mushroom stems can be used to produce a meat analogue, microbial protein, or even extract carotenoids from the spores. Additionally, the fermented product can be processed into a minced meat format, as shown in **Figure 2C**.

**Figure 2:**
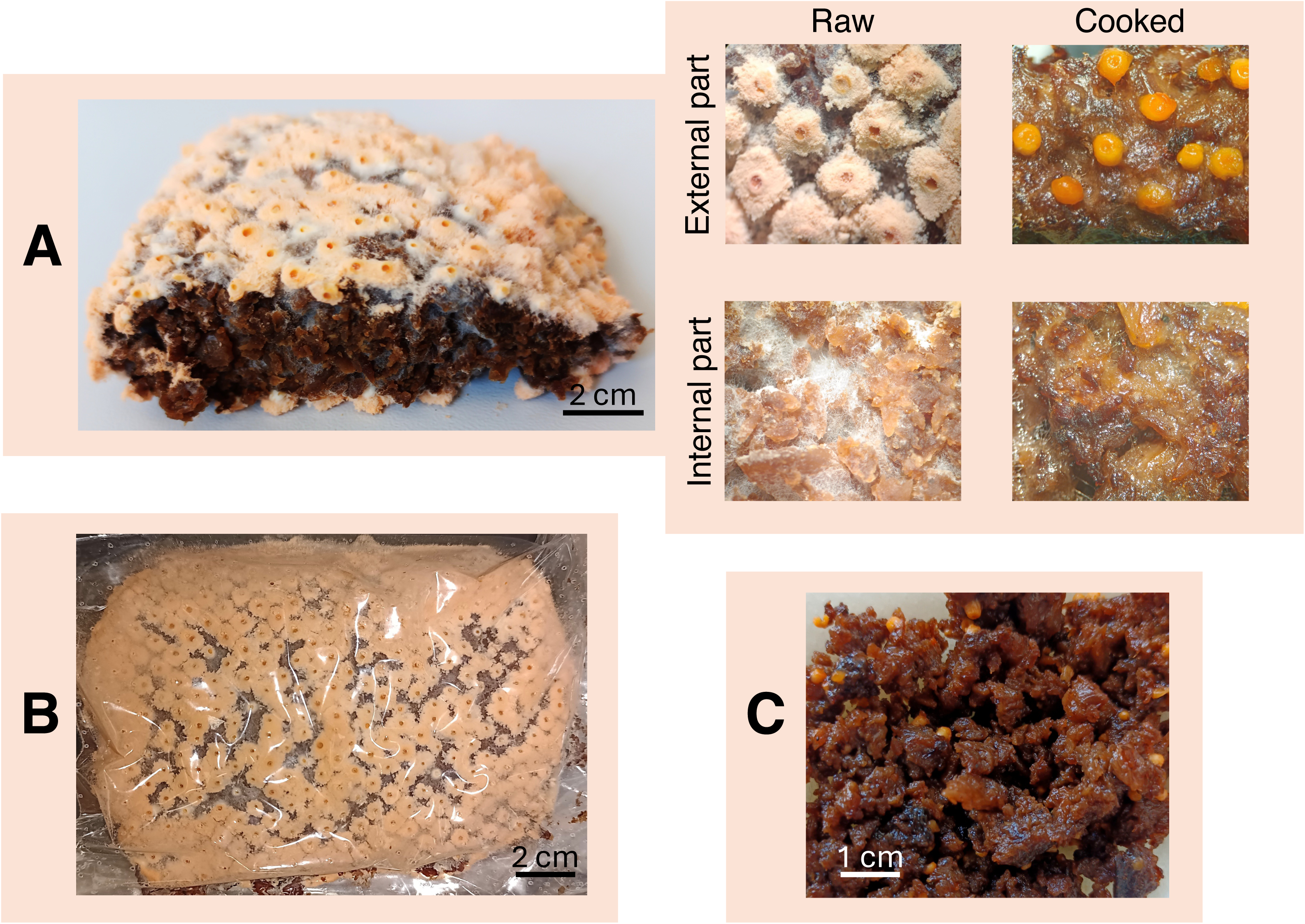
Appearance of 100 g of oyster mushroom stems fermented by *Neurospora crassa* under solid-state conditions after 48 h of incubation at 28°C in perforated polyethylene bags. This method is commonly used for industrial tempeh production. The image shows the fermented product cut in half (A), its appearance before and after cooking, its aspect during fermentation inside the plastic bags (B), and an example of the final product in a ground meat-like format (C).

### 3.2 Color analysis

In the color analysis, significant differences were observed between the cooked and uncooked *N. crassa* patties, as well as in comparison to the control samples, tempeh and meat patties. Regarding lightness (L), no significant differences were found between the top and bottom of the mushroom patty, but clear differences were noted when compared to the cooked meat (*p* < 0.001) (**Table 1**). Furthermore, a distinct change in lightness was observed between the cooked and uncooked mushroom patties, which is attributed to the loss of opacity in the hyphal structure during cooking (*p* < 0.001).

**Table 1.**
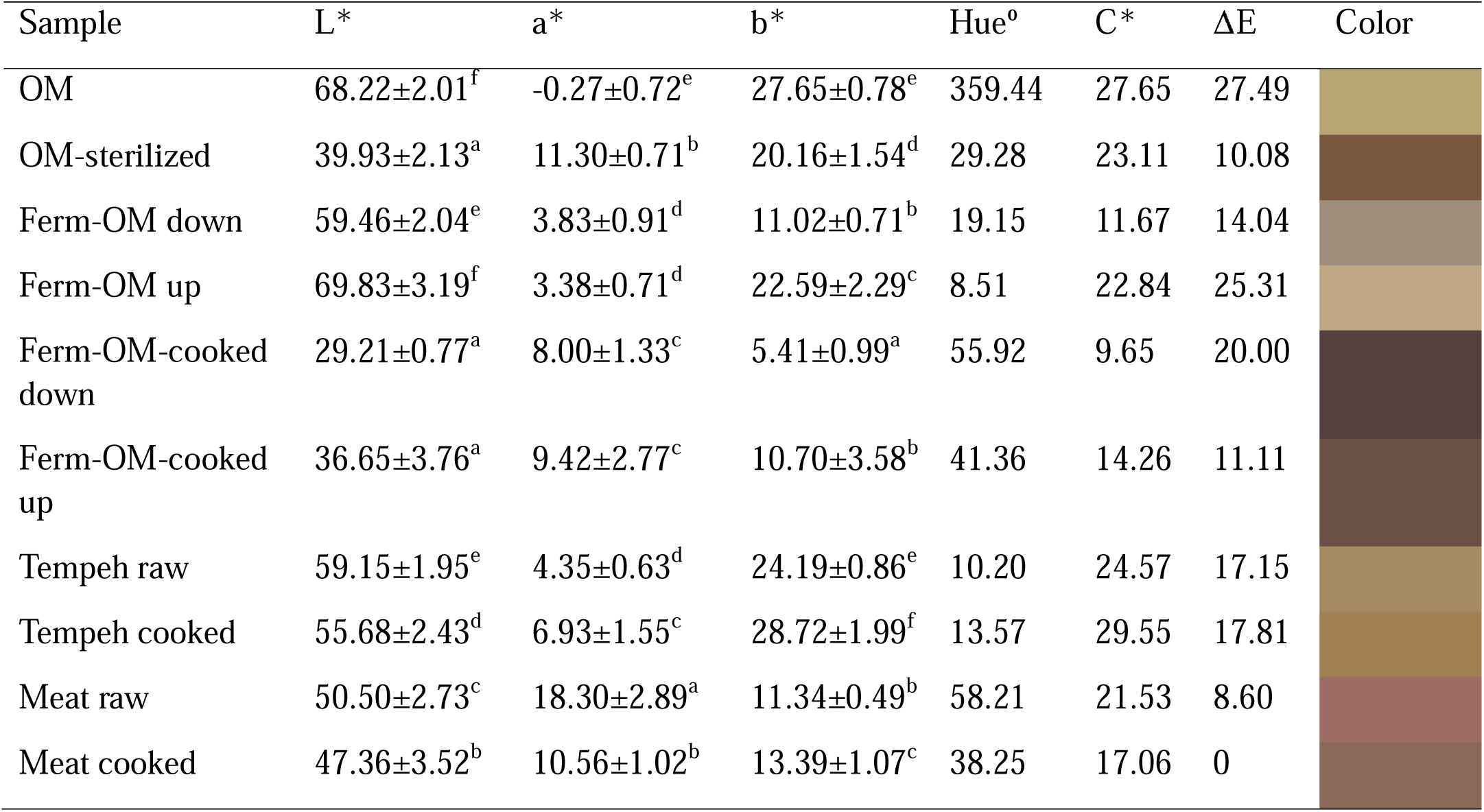
Color analysis of the different samples using the L, a, b color system. Values are expressed as mean ± standard deviation (n = 3). Statistical differences among samples were evaluated by one-way ANOVA followed by Tukey’s post hoc test (*p* < 0.05). Hue° (°Hue), Chroma (C*), and ΔE*ab were calculated from the L*, a*, b* values, with ΔE*ab indicating the color difference relative to Meat cooked.

| Sample | L* | a* | b* | Hue° | C* | $\Delta E$ | Color |
| --- | --- | --- | --- | --- | --- | --- | --- |
| OM | 68.22±2.01<sup>f</sup> | -0.27±0.72<sup>e</sup> | 27.65±0.78<sup>e</sup> | 359.44 | 27.65 | 27.49 | |
| OM-sterilized | 39.93±2.13<sup>a</sup> | 11.30±0.71<sup>b</sup> | 20.16±1.54<sup>d</sup> | 29.28 | 23.11 | 10.08 | |
| Ferm-OM down | 59.46±2.04<sup>e</sup> | 3.83±0.91<sup>d</sup> | 11.02±0.71<sup>b</sup> | 19.15 | 11.67 | 14.04 | |
| Ferm-OM up | 69.83±3.19<sup>f</sup> | 3.38±0.71<sup>d</sup> | 22.59±2.29<sup>c</sup> | 8.51 | 22.84 | 25.31 | |
| Ferm-OM-cooked down | 29.21±0.77<sup>a</sup> | 8.00±1.33<sup>c</sup> | 5.41±0.99<sup>a</sup> | 55.92 | 9.65 | 20.00 | |
| Ferm-OM-cooked up | 36.65±3.76<sup>a</sup> | 9.42±2.77<sup>c</sup> | 10.70±3.58<sup>b</sup> | 41.36 | 14.26 | 11.11 | |
| Tempeh raw | 59.15±1.95<sup>e</sup> | 4.35±0.63<sup>d</sup> | 24.19±0.86<sup>e</sup> | 10.20 | 24.57 | 17.15 | |
| Tempeh cooked | 55.68±2.43<sup>d</sup> | 6.93±1.55<sup>c</sup> | 28.72±1.99<sup>f</sup> | 13.57 | 29.55 | 17.81 | |
| Meat raw | 50.50±2.73<sup>c</sup> | 18.30±2.89<sup>a</sup> | 11.34±0.49<sup>b</sup> | 58.21 | 21.53 | 8.60 | |
| Meat cooked | 47.36±3.52<sup>b</sup> | 10.56±1.02<sup>b</sup> | 13.39±1.07<sup>c</sup> | 38.25 | 17.06 | 0 | |

For the red-green axis (a*), meat raw exhibited the highest value (18.30 ± 2.89^a^), significantly higher than all other samples. OM-sterilized and meat cooked also showed relatively high a* values, while OM (-0.27 ± 0.72 ^e^) and Ferm-OM up (3.38 ± 0.71 ^d^) had the lowest, indicating a much greener hue. In OM and Ferm-OM, a slight increase in red intensity was observed upon cooking or sterilization. On the yellow-blue axis (b*), tempeh cooked and OM showed the highest yellow values, while Ferm-OM-cooked down had the lowest, indicating a much bluer or less yellow hue.

An overall view of the results (**Figure 3A**) shows that, in terms of color attributes, Ferm-OM-cooked is positioned close to the cooked meat. According to **Table 1**, the hue of Ferm-OM-cooked is very similar to that of meat and noticeably closer than tempeh. The chroma of the upper portion of Ferm-OM-cooked is also comparable to cooked meat, and the color difference (ΔE = 11.11) indicates a perceptible but moderate distinction. While Ferm-OM-cooked approximates the visual appearance of meat, it is not identical. In contrast, raw Ferm-OM is closer to tempeh in both hue and chroma due to the whitish color of the hyphae, which is reflected in its lower saturation and distinct hue.

**Figure 3:**
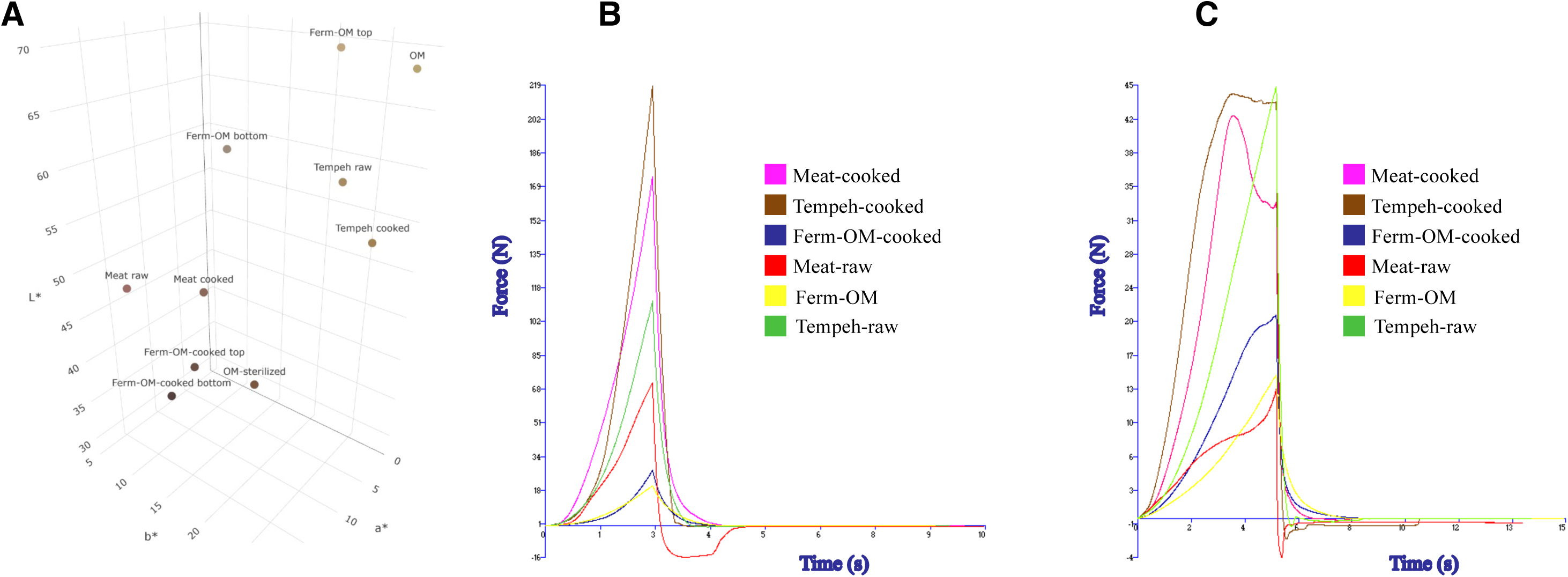
Color, compression and penetration analysis of meat analogues based on oyster mushroom, tempeh and meat patties (A) Graphical representation of color using b*, a* and L* values as x, y and z axes respectively. Raw oyster mushroom powder (OM), sterilized powder (OM-sterilized), solid-state fermentation of oyster mushroom (Ferm-OM), its cooked version (Ferm-OM-cooked), tempeh and meat cooked and uncooked are analyzed. As for texture analysis, the compression test (B) and the penetration test (C) are shown.

### 3.3 Texture analysis

The texture analysis revealed high heterogeneity among the samples. In the compression test, cooked tempeh presented the highest compressive strength, surpassing even cooked meat, suggesting a compact structure (**Figure 3B**). In contrast, Ferm-OM and Ferm-OM-cooked showed significantly lower values of compressive strength and area (*p* < 0.001), confirming its softer and more deformable character compared to meat and tempeh (**Supplementary Table 1**). Regarding elastic recovery after compression, this was appreciable only in the raw meat, while in the rest of the samples the behavior was predominantly plastic, evidenced by negative forces close to zero and a reduced negative displacement after unloading.

A similar trend was observed in the penetration test, where the fermented mushroom consistently showed lower resistance than cooked meat (*p* < 0.001) (**Figure 3C**). Cooking increased the firmness of all samples as evidenced by increases in positive peak strength, positive area, and slope to peak (**Supplementary Table 2**). In particular, cooked tempeh showed the greatest resistance to penetration, while cooked meat also showed significant toughening compared to its raw counterpart. A sudden drop in force around the 5-second mark is observed in all cases, indicating structural failure or rupture (**Figure 3C**). In terms of behavior after probe removal, most samples showed a plastic pattern with little elastic recovery, a phenomenon that was more pronounced in cooked samples.

### 3.4 Microstructure

Samples of OM, OM-sterilized, Ferm-OM and Ferm-OM-cooked were analyzed in the FESEM. The results showed significant differences in the microstructure of the four samples (**Figure 4**). As for OM, it presents a topology composed of short fibers and granules of irregular size (**Figure 4A**). Once the substrate is autoclaved the topology becomes smooth, although striations and a network of holes can be seen due to the fusion of the oyster mushroom fibers (**Figure 4B**).

**Figure 4:**
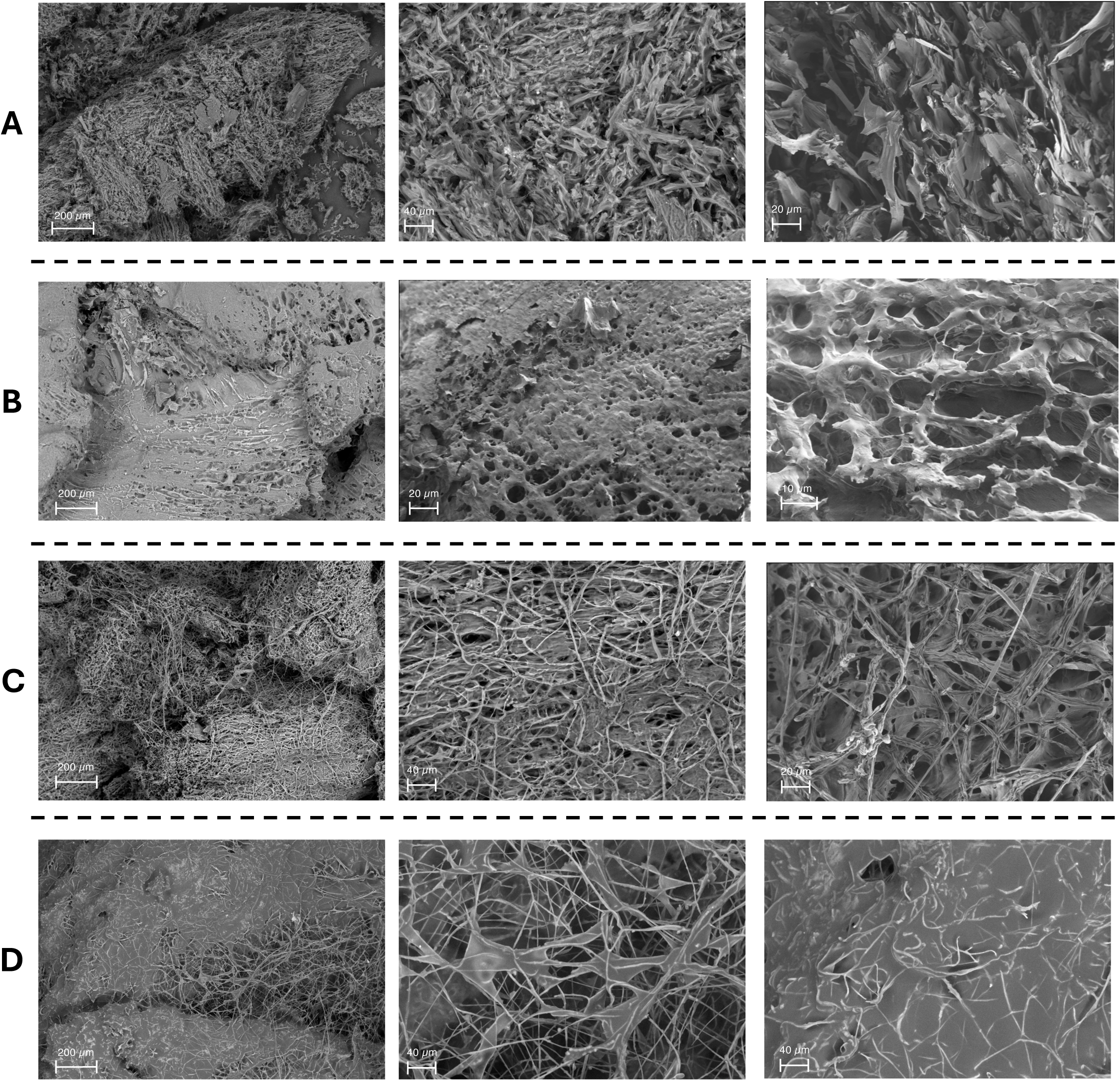
Field emission scanning electron microscopy (FESEM) micrographs of mushroom-based products. (A) Raw oyster mushroom stem powder (OM). (B) Sterilized oyster mushroom stem powder (OM-sterilized). (C) Solid-state fermentation of oyster mushroom stem waste with *Neurospora crassa* (Ferm-OM). (D) Stir fried Ferm-OM (Ferm-OM-cooked).

The microstructure changes drastically after fermentation (**Figure 4C**). At this time, the hyphae of *N. crassa* completely colonize the substrate and form a network that envelops the substrate granules and connects them to each other. The hyphae between granules explain why this composite does not crumble and why the product adopts a firm conformation. A closer view shows that the holes in the autoclaved substrate could facilitate the adhesion of *N. crassa* hyphae. Once the product is cooked (**Figure 4D**), the topology of the product becomes more compact. The hyphae surrounding the granules fuse with the substrate and a smooth, hole-free topology. As for the hyphae connecting the granules, these are still present, although much of them have fused, increasing the thickness of the filament. This microstructure explains that, once the product is cooked, it remains firm and cohesive.

### 3.5 Technofunctional properties

The water retention, water absorption, and soluble content of lyophilized samples varied depending on the processing conditions (**Table 2**). OM exhibited a water retention capacity of 6.12±0.69 g/g and a water absorption capacity of 6.79±0.51 g/g, with 287.60±20 mg/g of soluble compounds. Sterilization significantly increased water retention (8.33±0.66 g/g) and absorption (8.74±0.44 g/g), while slightly reducing soluble compounds (258.23±25 mg/g), likely due to heat-induced modifications in the structural matrix and leaching of these components.

**Table 2.** Technofunctional properties of lyophilised samples. Values are expressed as mean ± standard deviation (n = 3). Statistical differences among samples were evaluated by one-way ANOVA followed by Tukey’s post hoc test (*p* < 0.05).

| Sample | H<sub>2</sub>O retention (g/g) | H<sub>2</sub>O absorption (g/g) | Solubility (mg/g) | Oil retention (g/g) |
| --- | --- | --- | --- | --- |
| OM | 6.12±0.69<sup>b</sup> | 6.79±0.51<sup>bc</sup> | 287.60±20<sup>ab</sup> | 12.30±0.70<sup>a</sup> |
| OM-sterilized | 8.33±0.66<sup>a</sup> | 8.74±0.44<sup>a</sup> | 258.23±25<sup>b</sup> | 4.70±0.17<sup>c</sup> |
| Ferm-OM | 6.50±0.28<sup>b</sup> | 7.56±0.10<sup>ab</sup> | 313.50±9<sup>a</sup> | 6.00±0.23<sup>b</sup> |
| Ferm-OM-cooked | 5.35±0.56<sup>b</sup> | 5.89±0.51<sup>c</sup> | 256.61±5<sup>b</sup> | 2.57±0.10<sup>d</sup> |

Ferm-OM showed lower water retention (6.50±0.28 g/g) and absorption (7.56±0.10 g/g) than Ferm-OM-cooked but showed the highest soluble compounds content (313.50±9 mg/g), suggesting enhanced solubilization of compounds due to microbial activity. However, cooking the patty reduced water retention (5.35±0.56 g/g) and absorption (5.89±0.51 g/g), alongside a marked decrease in soluble compounds (256.61±5 mg/g), indicating structural modifications that may have led to reduced hydration properties and solubilization losses.

Oil retention capacity varied significantly among the different samples (**Table 2**). OM exhibited the highest oil retention, whereas sterilization of the substrate by high-temperature autoclaving led to a marked reduction in this capacity. Following fermentation, oil retention was higher than in the autoclaved substrate; however, this trend reversed after cooking. The Ferm-OM-cooked sample showed the lowest oil retention values, although this effect is partially masked by the fact that approximately 5 mL of oil was used per burger during the cooking process.

### 3.6 Nutritional composition

The differences in nutritional composition between the oyster mushroom (OM) and the fermented product (Ferm-OM) are notable. First, a reduction in protein content is observed, which could be attributed to the consumption of nitrogenous compounds by *N. crassa* during fermentation (*p* < 0.05) (**Table 3**). Therefore, there is a decrease in the total amino acid content, although their relative profile is maintained in similar proportions (**Table 4**). The non-essential amino acids glutamic acid and aspartic acid, which are the most abundant and contribute to the umami taste, significantly decrease after fermentation (*p* < 0.05). Regarding the essential amino acids, a decrease in phenylalanine and methionine is observed (*p* < 0.05).

**Table 3.** Proximate composition of OM and Ferm-OM (g/100g dw). Values are expressed as mean ± standard deviation (n = 3). Statistical differences between groups were evaluated using a two-sample t-test (*p* < 0.05).

| Component | OM | Ferm-OM | *p* value |
| --- | --- | --- | --- |
| Protein | 16.6±1.5 | 11.5±1.0 | < 0.05 |
| Fat | 2.1±0.3 | 2.6±0.4 | ns |
| Total fibre | 51.5±7.3 | 67.3±8.4 | < 0.05 |
| Total sugars | 2.3±0.4 | 3.1±0.5 | ns |
| Insoluble carbohydrates | 26.0±2.6 | 23.8±2.3 | ns |
| Sodium | 0.012±0.0002 | 0.028±0.004 | < 0.01 |
| NaCl | 0.030±0.0005 | 0.071±0.009 | < 0.001 |
| Moisture (%) | 79.04±1.38 | 78.72±0.69 | ns |
| Energy dw (kcal/100g) | 197.50 | 216.40 |  |
| Energy ww (kcal/100g) | 41.40 | 46.05 |  |

**Table 4.** Amino acid composition (%).Values are expressed as mean ± standard deviation (n = 3). Statistical differences between groups were evaluated using a two-sample t-test (*p* < 0.05).

| Amino acid | OM | Ferm-OM | *p* value |
| --- | --- | --- | --- |
| Aspartic acid | 1.06±0.08 | 0.70±0.09 | < 0.05 |
| Glutamic acid | 1.26±0.10 | 0.89±0.11 | < 0.05 |
| Alanine | 0.58±0.09 | 0.42±0.08 | ns |
| Arginine | 0.45±0.08 | 0.29±0.06 | ns |
| Cystine + Cysteine | 0.14±0.03 | 0.15±0.03 | ns |
| Phenylalanine | 0.38±0.06 | 0.23±0.04 | < 0.05 |
| Glycine | 0.44±0.06 | 0.39±0.07 | ns |
| Histidine | 0.20±0.02 | 0.14±0.03 | ns |
| Isoleucine | 0.48±0.05 | 0.47±0.06 | ns |
| Leucine | 0.86±0.11 | 0.83±0.10 | ns |
| Lysine | 0.53±0.07 | 0.37±0.07 | ns |
| Methionine | 0.14±0.02 | 0.09±0.01 | < 0.05 |
| Proline | 0.38±0.07 | 0.28±0.05 | ns |
| Serine | 0.46±0.07 | 0.33±0.06 | ns |
| Tyrosine | 0.29±0.05 | 0.16±0.03 | < 0.05 |
| Threonine | 0.44±0.04 | 0.34±0.06 | ns |
| Tryptophan | 0.15±0.03 | 0.10±0.02 | ns |
| Valine | 0.46±0.06 | 0.37±0.07 | ns |

Moreover, fermentation provides improvements in nutritional values, mainly through an increase in fiber and fatty acid content. It is important to note that *N. crassa* is a natural source of chitin, β-glucans and other structural polysaccharides characteristic of fungi (Bartholomai et al., 2022). Fermentation enriches the substrate with additional fungal biomass, thus increasing the proportion of cell walls and membranes, resulting in an increase in total fiber and lipid content.

Regarding the lipid fraction, both products have a profile dominated by unsaturated fatty acids, with a high content of linoleic acid (C18:2n6) (**Table 5**). In addition, fermentation significantly increases the content of omega-3 fatty acids, in particular alpha-linolenic acid (C18:3n3), whose level in Ferm-OM is more than 15 times that of the unfermented oyster mushroom (*p* < 0.001). This enrichment in essential fatty acids contributes to improving the overall lipid profile of the product, adding functional value in terms of cardiovascular health and anti-inflammatory properties (Djuricic & Calder, 2021).

**Table 5.** Fatty acid composition (g/100 g dw). Values are expressed as mean ± standard deviation (n = 3). Statistical differences between groups were evaluated using a two-sample t-test (*p* < 0.05).

| Fatty acid | OM | Ferm-OM | *p* value |
| --- | --- | --- | --- |
| Butyric C4:0 | <0.002 | <0.002 |  |
| Caproic C6:0 | <0.002 | <0.002 |  |
| Caprylic C8:0 | <0.002 | <0.002 |  |
| Capric C10:0 | <0.002 | <0.002 |  |
| Undecanoic C11:0 | <0.002 | <0.002 |  |
| Lauric C12:0 | <0.002 | <0.002 |  |
| Tridecanoic C13:0 | <0.002 | <0.002 |  |
| Myristic C14:0 | 0.003±0.001 | <0.002 | |
| Myristoleic C14:1 | <0.002 | <0.002 |  |
| Pentadecanoic C15:1 | 0.024±0.003 | 0.031±0.004 | < 0.05 |
| Pentadecenoic C15:1 | <0.002 | <0.002 |  |
| Palmitic C16:0 | 0.225±0.005 | 0.514±0.007 | < 0.001 |
| Palmitoleic C16:1 | 0.007±0.001 | 0.022±0.003 | < 0.001 |
| Margaric C17:0 | 0.004±0.001 | <0.002 | |
| Margaroleic C17:1 | 0.002±0.001 | <0.002 | |
| Stearic C18:0 | 0.052±0.002 | 0.216±0.003 | < 0.001 |
| Trans-Oleic C18:1n9 | 0.002±0.001 | <0.002 | |
| Oleic C18:1n9 | 0.191±0.002 | 0.246±0.003 | < 0.001 |
| Trans-Linoleic C18:2n6 | 0.002±0.001 | <0.002 | |
| Linoleic C18:2n6 | 1.551±0.020 | 1.445±0.018 | < 0.01 |
| Gamma-Linolenic C18:3n6 | <0.002 | <0.002 |  |
| Trans-Linolenic C18:3 | <0.002 | <0.002 |  |
| Arachidic C20:0 | 0.002±0.001 | <0.002 | |
| Alpha-Linolenic C18:3n3 | 0.004±0.001 | 0.071±0.001 | < 0.001 |
| Gadoleic C20:1n9 | 0.002±0.001 | <0.002 | |
| Heneicosanoic C21:0 | <0.002 | <0.002 |  |
| Eicosadienoic C20:2n6 | 0.003±0.001 | <0.002 |  |
| Dihomo-Gamma-Linolenic C20:3n6 | <0.002 | <0.002 |  |
| Behenic C22:0 | 0.002±0.001 | <0.002 |  |
| Eicosatrienoic C20:3n3 | <0.002 | <0.002 |  |
| Arachidonic C20:4n6 | <0.002 | <0.002 |  |
| Erucic C22:1n9 | 0.002±0.001 | <0.002 |  |
| Tricosanoic C23:0 | <0.002 | <0.002 |  |
| Docosadienoic C22:2n6 | <0.002 | <0.002 |  |
| Eicosapentaenoic acid C20:5n3 | <0.002 | <0.002 |  |
| Lignoceric C24:0 | 0.004±0.001 | <0.002 |  |
| Nervonic C24:1n9 | 0.012±0.003 | <0.002 |  |
| Docosahexaenoic Acid C22:6n3 | <0.002 | <0.002 |  |
| Saturated fatty acids | 0.316±0.002 | 0.762±0.001 | < 0.001 |
| Cis-monounsaturated fatty acids | 0.216±0.025 | 0.268±0.044 | ns |
| Cis, cis-polyunsaturated fatty acids | 1.557±0.025 | 1.516±0.020 | ns |
| Total trans unsaturated fatty acids | 0.013±0.002 | <0.002 |  |
| Omega-3 fatty acids | 0.004±0.001 | 0.071±0.001 | < 0.001 |
| Omega-6 fatty acids | 1.553±0.022 | 1.445±0.019 | < 0.01 |
| Omega-9 fatty acids | 0.207±0.005 | 0.246±0.003 | < 0.001 |

### 3.7 Prebiotic potential

In this study, the differential growth of prebiotic bacteria and *E. coli* was used to evaluate the effect of fermentation and in vitro digestion on the prebiotic potential of the samples. Growth curves revealed that digestion altered bacterial behavior, generally promoting a shift towards the utilization of more complex substrates such as Ferm-OM, while reducing growth on simple substrates such as fructooligosaccharides (**Figure 5**). This effect was particularly evident in *B. bifidum*, *B. longum*, *L. lactis*, and *S. salivarius*. In contrast, for the pathogenic bacterium *E. coli*, growth on OM, Ferm-OM, and Ferm-OM-cooked decreased after digestion.

**Figure 5.**
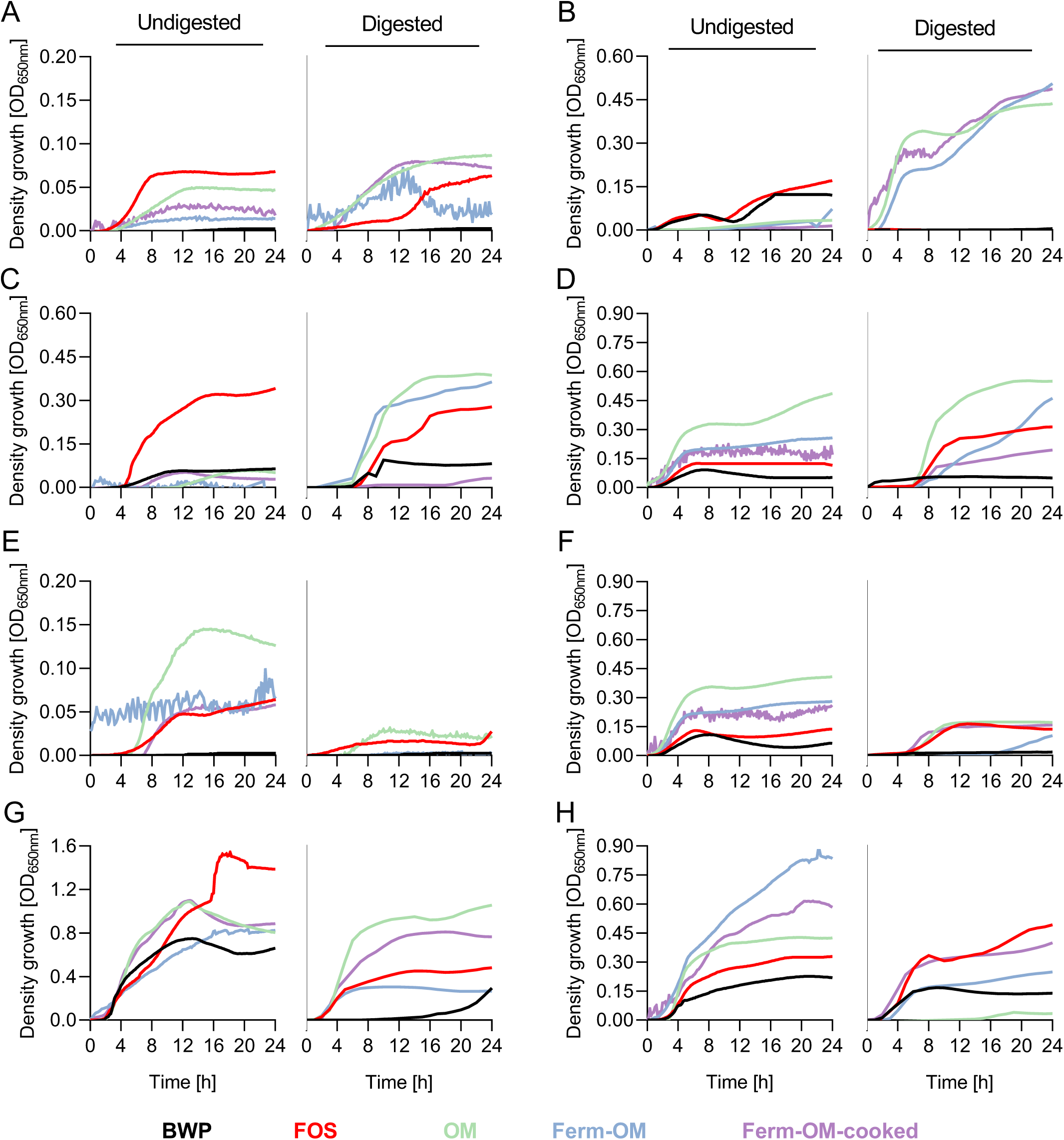
Growth of representative beneficial bacteria stimulated by undigested or digested mushroom-based products: raw oyster mushroom stem powder (OM), Solid-state fermentation of OM with *Neurospora crassa* (Ferm-OM), and stir fried Ferm-OM (Ferm-OM-cooked). Beneficial bacteria included (A) *Bifidobacterium bifidum*, (B) *B. longum*, (C) *Lactococcus lactis*, (D) *Lactobacillus brevis*, (E) *L. plantarum*, (F) *Pediococcus acidilactici*, and (G) *Streptococcus salivarius*, while (H) *Escherichia coli* was included as a pathogenic representative. Buffered peptone water (BWP) and fructooligosaccharides (FOS) were included as negative and positive controls, respectively. All substrates represented 0.2% w/v final volume. Bacterial growth was expressed as mean optical density values measured at 650 nm (OD_650nm_).

In the undigested samples, *L. lactis* and *S. salivarius* showed significantly higher final and maximum growth in the presence of FOS (*p* < 0.001) (**Figure 6**). After substrate digestion, *L. lactis* exhibited greater growth with OM and Ferm-OM, whereas *S. salivarius* grew more with OM and Ferm-OM-cooked. *B. longum* displayed selective growth in the presence of OM, Ferm-OM, and Ferm-OM-cooked only after digestion (*p* < 0.001). Some bacteria, such as *L. brevis*, metabolized the substrates similarly regardless of digestion, while *P. acidilactici* grew more abundantly with undigested substrates. In contrast, *B. bifidum* and *L. plantarum* showed poor growth under any condition. Finally, *E. coli* exhibited significant growth in the presence of OM, Ferm-OM, and Ferm-OM-cooked in undigested samples (*p* < 0.001); however, after digestion, maximum growth with OM and Ferm-OM-cooked decreased.

**Figure 6.**
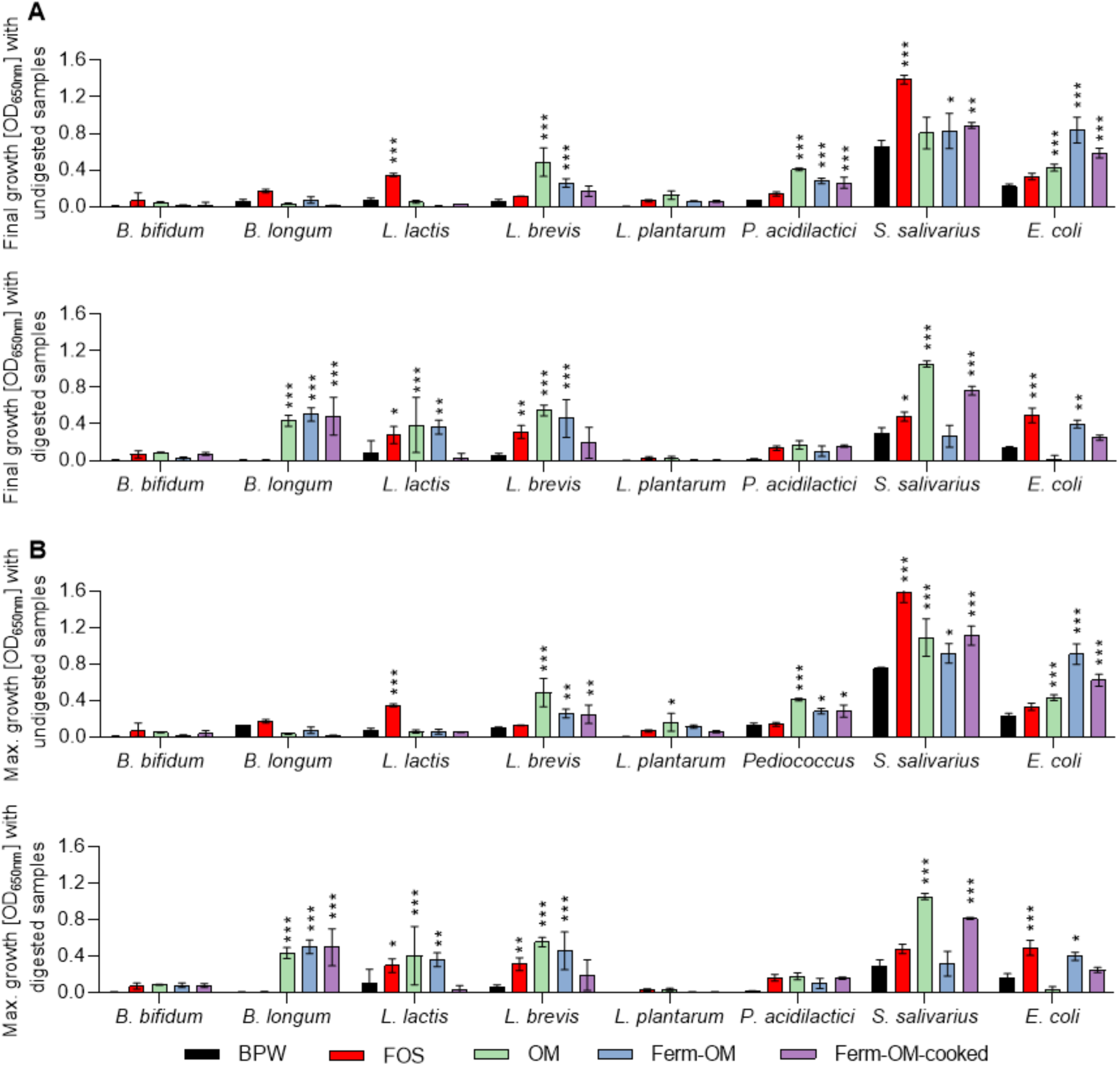
(A) Final growth and (B) maximum growth of representative beneficial bacteria stimulated by undigested or digested mushroom-based products: raw oyster mushroom stem powder (OM), Solid-state fermentation of OM with *Neurospora crassa* (Ferm-OM), and stir fried Ferm-OM (Ferm-OM-cooked). Buffered peptone water (BWP) and fructooligosaccharides (FOS) were included as negative and positive controls, respectively (\*\*\**p*<0.001, \*\**p*<0.01, \**p*<0.05, ns *p*>0.05; according to two-way ANOVA with Dunnett’s post hoc test). All substrates represented 0.2% w/v of final volume. Bacterial growth was expressed as mean optical density values (n=3) ± standard deviation, measured at 650 nm (OD_650nm_).

In this study, the PAS was calculated by comparing the growth of prebiotic bacteria with that of the pathogenic bacterium *E. coli* (**Figure 7A**). Overall, the undigested samples showed the lowest PAS values, particularly Ferm-OM and Ferm-OM-cooked. Among the few positive values were OM with *B. bifidum* (*p* < 0.01), *L. brevis* (*p* < 0.01), and *P. acidilactici* (*p* < 0.05). After digestion, a stronger affinity of prebiotic bacteria for the substrates was observed compared to *E. coli*. Notably, *B. longum* exhibited the most positive values with Ferm-OM and Ferm-OM-cooked. For OM, PAS values were positive across all tested bacteria. In contrast, Ferm-OM-cooked, although showing relatively high values in most cases, yielded positive PAS only with *B. bifidum*, *B. longum*, *P. acidilactici*, and *S. salivarius*.

**Figure 7.**
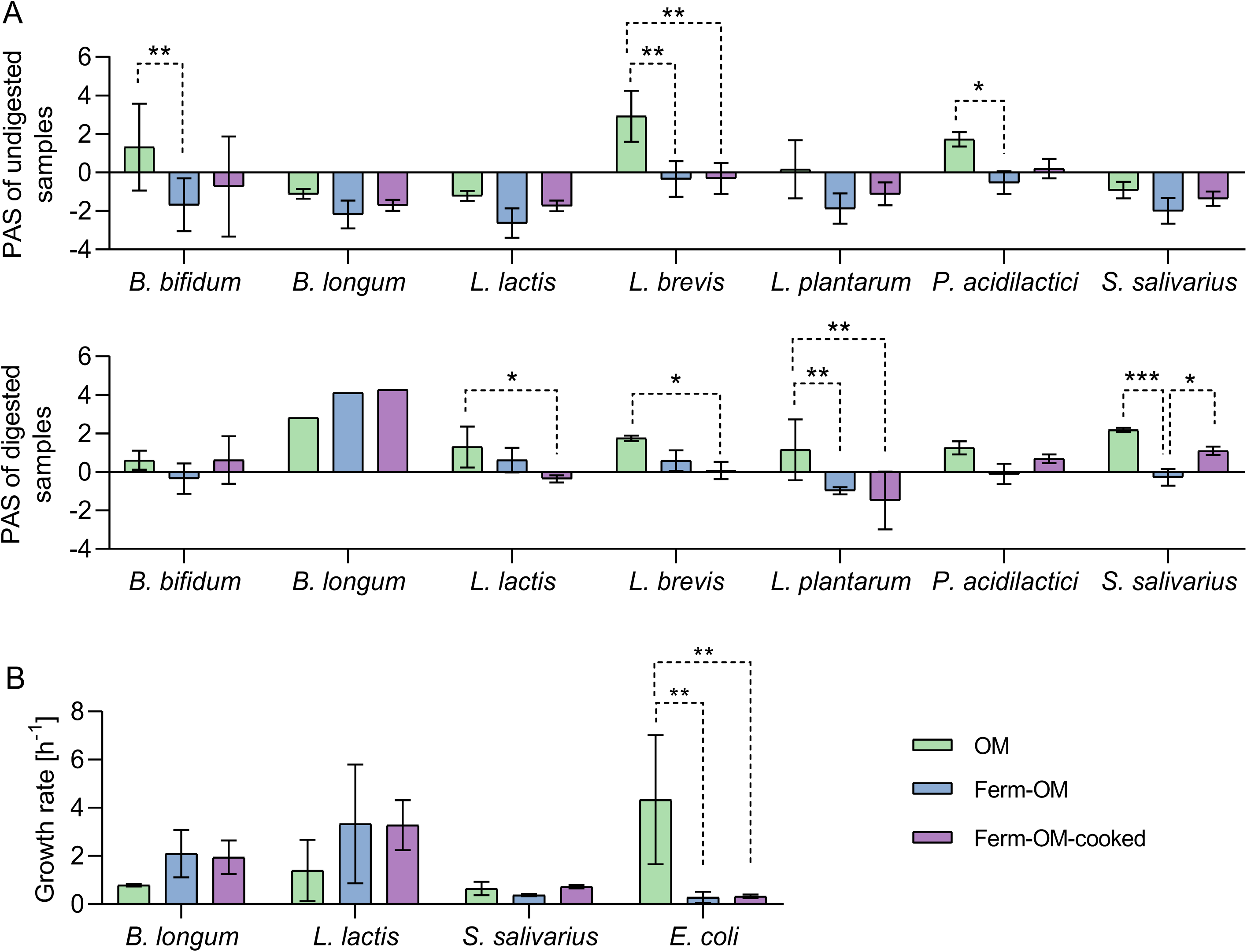
(A) Prebiotic activity score (PAS) of undigested or digested mushroom-based products: raw oyster mushroom stem powder (OM), solid-state fermentation of OM with *Neurospora crassa* (Ferm-OM), and stir fried Ferm-OM (Ferm-OM-cooked), in beneficial bacteria. (B) Growth rate calculated from growth curves of bacteria with positive PAS, exposed to digested samples. Samples were compared with each other (\*\*\**p*<0.001, \*\**p*<0.01, \**p*<0.05; according to two-way ANOVA with Tukey’s post hoc test). Data was expressed as mean values (n=3) ± standard deviation.

### 3.8 Microbiological analysis

The results of the microbiological analysis showed that no pathogenic bacteria have been detected in the final product, which reinforces the safety of SSF as a method to produce novel foods. As for the bacteria tested, aerobic mesophilic bacteria, total coliforms, *Escherichia coli*, *Listeria monocytogenes* and coagulase-positive *Staphylococcus* were below the detection limit (<10.00 CFU/g). For the last test, *Salmonella* spp. was not detected in 25 g of the sample.

## 4. Discussion

Fungi of the genus *Neurospora* have shown great potential for the conversion of agro-industrial wastes into novel food products or biomass (Maini Rekdal et al., 2024; Toghiani et al., 2025). In this study, we successfully developed a meat analog using oyster mushroom stipes fermented under solid-state conditions by a *N. crassa* strain isolated in Albacete, Spain. This by-product is particularly promising due to its high fiber content, although it has not been previously explored as a substrate for SSF. Through SSF, we enriched *P. ostreatus* with 15.8% more total fibre, resulting in a product not only richer in functional properties but also potentially beneficial for gut health, offering an enhanced alternative to oyster mushrooms alone.

In SSF, the composition of the substrate directly influences the composition of the final product. The oyster mushroom is characterized by a very low protein content and a majority share of carbohydrates (Araújo-Rodrigues et al., 2025; Bermúdez-Gómez et al., 2024). The fermented product, therefore, retains much of the fiber even though some of the protein has been metabolized by *N. crassa*. This large amount of fiber differentiates this product from the other meat analogues that have been developed or from fermented products such as tempeh, which is richer in protein due to the fermentation of soybeans (Thulesen et al., 2025). The presence of nitrogen seems to be a limiting factor for protein production by *N. crassa*, so using variable proportions of other fungi such as *Agaricus bisporus* (≈15 and 40 g/100g dw in stem and cap, respectively) (Alnoumani et al., 2017; Bermúdez-Gómez et al., 2024; Krishnamoorthi et al., 2022) or *Lentinula edodes* (≈15 g/100g dw in the cap) (Zeng et al., 2024) might considerably increase the protein proportion of the fermented product.

Despite not having a particularly high protein content, its unsaturated fatty acids, polysaccharides, the presence of essential amino acids, umami and the good digestibility of oyster mushroom and *Neurospora* proteins increase the nutritional quality of this product. Fermentation with filamentous fungi, in short, may increase the protein ratio and breaks down complex carbohydrates and proteins, resulting in a more digestible product (Li et al., 2019). In addition, fermentation between strains of filamentous fungi, which degrade complex polysaccharides, with bacteria such as *Lactiplantibacillus plantarum*, or the supplementation with ammonium salts can significantly increase the protein content of the final product (Canedo et al., 2016; Feng et al., 2024). Despite ongoing efforts to increase protein content in meat analogues, this study introduces an innovative alternative: products that visually resemble meat but have a radically different nutritional composition. The high fibre content of this product offers a promising strategy to help meet the recommended daily intake of 25 grams/day, supporting bowel function and well-being (EFSA Panel on Dietetic Products, Nutrition, and Allergies (NDA), 2010). Moreover, the final fermented product, due to the high presence of fibre, has a much higher water-holding capacity than conventional meat, which could therefore increase the feeling of satiety in consumers (Song et al., 2025).

The presence of dietary fiber, in addition to supporting intestinal transit, has been associated with shifts in the gut microbiota that may influence certain diseases (Gill et al., 2021). In this study, it was observed that after *in vitro* digestion, substrates generally became more accessible to beneficial bacteria. This occurs because prebiotic polysaccharides resist digestion in the upper gastrointestinal tract and reach the colon, where they are fermented by these bacteria, whereas simpler molecules are absorbed in the small intestine (Liu et al., 2025). The fermentation of OM by *N. crassa* results in a product richer in fiber, but this does not necessarily translate into increased prebiotic activity. In contrast, when the product is cooked, the fiber increases its bioaccessibility and can be more extensively metabolized by probiotic bacteria. This may be due to the fact that while *N. crassa* consumes part of the nutrients to increase insoluble fiber, cooking alters the structure of these polysaccharides, modifying and promoting their solubility and fermentability (Fang et al., 2023; Nowak et al., 2025). Moreover, *in vitro* digestion of OM, Ferm-OM, and Ferm-OM-cooked removes easily assimilable molecules while leaving fungal β-glucans largely intact and available for specific fermentation by probiotic bacteria, thereby increasing their selectivity against *E. coli* (Li et al., 2024).

Another aspect to take into consideration in the formulation of Ferm-OM is the texture of the final product. While the fermented product has a cohesive structure, it has less toughness than conventional meat patties or tempeh. Ferm-OM-cooked, although significantly softer, showed a slight increase in strength after cooking, suggesting a slight structural reorganization of the matrix during heat treatment. However, its texture remained soft, with low strength values at 5 mm and inclination, which could translate into easier chewing and higher palatability. In the case of Ferm-OM, higher fermentation times, such as 72 h, could result in products with more compact textures. In addition, increasing the fermentation time in SSF has been shown to further increase the microbial protein content (Toghiani et al., 2025). Despite this, it is not recommended to exceed the fermentation time because fermentation produces free amino acids, some of which can produce a bitter taste (Utami et al., 2016).

The problem of exceeding the fermentation time is the appearance of a large number of spores that, despite being rich in carotenoids, affect the final appearance of the product and visually impair it if it claims to be a meat analog. In this study, the characteristic brownish tone, typical of certain burgers, appeared after substrate sterilization but diminished during fermentation, reappearing once the product was cooked. This reversible color behavior suggests that heat-induced reactions, such as Maillard or caramelization processes, are the main contributors to the color change. For pilot-scale-up, fermentation in food grade zip-lock plastic bags that allow gas exchange, similar to the process used for tempeh, could bring us closer to the production of compact blocks of *oncom* that are easier to handle and have a higher yield (Fernandez Castaneda et al., 2024). The relationship between cohesiveness, taste, safety and nutritional content is one of the most relevant aspects to take into account when commercializing this new food.

At the industrial level, SSF is a technique that makes it possible to produce food from agricultural by-products using a low amount of water, which avoids contamination with pathogens. In this study, we start from freeze-dried oyster mushrooms, but this process is costly and expensive, so the industry needs to work with fresh mushrooms due to the large volume of waste generated continuously throughout the year. This study also explores the opportunity of using wild strains for the production of fermented foods, as well as their possible subsequent pilot scaling up. There is a great deal of fungal diversity to be explored at our fingertips. *Neurospora* strains are ideal for fermenting organic matter rich in complex polysaccharides, as this fungus feeds on decayed wood. The symbiosis between traditional Indonesian techniques to produce *oncom* and the exploration of new microorganisms merge in this work to promote the circular economy in edible mushroom production.

## 5. Conclusions

This study demonstrates the potential of *N. crassa* to upcycle *P. ostreatus* cultivation by-products through solid-state fermentation. The resulting product exhibits a meat-like microstructure supported by the fungal hyphal network, although with a softer texture and a distinct nutritional profile. Fermentation leads to protein consumption from the substrate, accompanied by a relative increase in fiber content. These characteristics promote a selective response from prebiotic bacteria, suggesting its potential application as a functional food ingredient. Future studies should focus on optimizing the formulation through the incorporation of additional substrates, assessing shelf life, and scaling up the process to enable industrial and commercial application.

## Supporting information

Manuscript

Manuscript

## Data availability

The data supporting our findings are available in the manuscript file or from the corresponding author upon request.

## Ethics statement

Not applicable.

## Author Contributions

**Pablo Navarro-Simarro:** Conceptualization, Data curation, Formal analysis, Investigation, Methodology, Writing – original draft, Writing – review & editing. **Bryan Moreno-Chamba:** Data curation, Formal analysis, Investigation, Writing – original draft. **Julio Salazar-Bermeo** Investigation, Writing – review & editing. **Lourdes Gómez-Gómez:** Investigation, Supervision, Validation, Writing – review & editing. **Ángela Rubio-Moraga:** Conceptualization, Investigation, Supervision, Validation, Writing – review & editing. **Alberto José López Jimenez:** Data curation, Formal analysis, Investigation, Writing – review & editing. **Nuria Martí-Bruña:** Investigation, Supervision, Validation, Writing – review & editing. **Oussama Ahrazem:** Conceptualization, Funding acquisition, Investigation, Project administration, Supervision, Validation, Writing – original draft .All authors reviewed, contributed to, and approved the final version of the manuscript.

## Acknowledgements

The authors would like to thank Setas Meli S.L. for mushrooms’ by-products supplies.

## Abbreviations

OM: oyster mushroom powder
OM-sterilized: autoclaved oyster mushroom powder
Ferm-OM: fermented oyster mushroom with *Neurospora crassa*
Ferm-OM-cooked: stir fried fermented oyster mushroom
SSF: solid-state fermentation

---

***Supplementary Figure 1:*** Solid-state fermentation of *Neurospora crassa* on various fungal substrates, such as oyster mushroom and shiitake stems (A). Additionally, filamentous fungi of industrial interest that can ferment oyster mushroom stems to varying degrees are shown in (B).

