## Supplementary material for "Solid-state fermentation of oyster mushroom by-products using *Neurospora crassa*: a sustainable approach for the development of novel meat analogues": Manuscript

| **Supplementary Table 1.** Texture profile parameters obtained from the compression assay for raw and cooked samples. Values are expressed as mean ± standard deviation (n = 3). Statistical differences among samples were evaluated by one-way ANOVA followed by Tukey’s post hoc test (*p* < 0.05). | | | | | | |
| --- | --- | --- | --- | --- | --- | --- |
|  | Sample | | | | | |
| Parameter | Raw meat | Cooked meat | Raw tempeh | Cooked tempeh | Ferm-OM | Ferm-OM-cooked |
| Positive force peak (N) | 60.59±10.42^bc^ | 174.04±2.41^a^ | 110.73±31.87^b^ | 206.80±41.12^a^ | 15.66±4.76^c^ | 29.81±3.08^c^ |
| Negative force peak (N) | -13.32±2.28^b^ | -0.01±0.01^a^ | -0.80±0.55^a^ | -0.66±0.05^a^ | -0.06±0.06^a^ | -0.03±0.01^a^ |
| Positive displacement peak (mm) | 5 | 5 | 5 | 5 | 5 | 5 |
| Negative displacement peak (mm) | -9.80±0.56^d^ | -9.51±0.05^d^ | -8.81±0.75^cd^ | -6.15±0.45^b^ | -7.20±1.04^bc^ | -3.92±0.78^a^ |
| Positive area (N·s) | 50.83±14.92^b^ | 184.73±0.58^a^ | 105.79±35.51^ab^ | 187.09±78.72^a^ | 18.80±6.81^b^ | 27.35±2.80^b^ |
| Negative area (N·s) | -14.90±8.02^b^ | 0 | -0.64±0.53^a^ | -0.23±0.04^a^ | -0.02±0.02^a^ | 0 |
| Area to positive peak (N·s) | 46.03±13.97^b^ | 143.30±0.27^a^ | 81.50±28.49^ab^ | 146.05±60.73^a^ | 11.95±4.52^b^ | 17.74±1.75^b^ |
| Slope to positive peak (N/s) | 24.18±4.16^cd^ | 69.33±0.96^ab^ | 44.11±12.70^bc^ | 82.91±22.12^a^ | 6.24±1.89^d^ | 11.87±1.22^d^ |
| Force at 5 mm (N) | 60.59±10.42^cd^ | 173.74±2.35^ab^ | 110.61±31.86^bc^ | 206.75±55.53^a^ | 15.65±4.76^d^ | 29.74±3.06^d^ |

| **Supplementary Table 2.** Texture profile parameters obtained from the penetration assay for raw and cooked samples. Values are expressed as mean ± standard deviation (n = 3). Statistical differences among samples were evaluated by one-way ANOVA followed by Tukey’s post hoc test (*p* < 0.05). | | | | | | |
| --- | --- | --- | --- | --- | --- | --- |
|  | Sample | | | | | |
| Parameter | Raw meat | Cooked meat | Raw tempeh | Cooked tempeh | Ferm-OM | Ferm-OM-cooked |
| Positive force peak (N) | 14.56±4.93^c^ | 43.21±1.14^a^ | 46.72±2.05^a^ | 43.41±0.83^a^ | 16.17±1.05^bc^ | 21.53±1.90^b^ |
| Negative force peak (N) | -3.95±1.58^b^ | -0.28±0.09^a^ | -2.10±1.89^ab^ | -2.05±1.03^ab^ | -0.08±0.03^a^ | -0.03±0.01^a^ |
| Positive displacement peak (mm) | 10 | 10 | 10 | 10 | 10 | 10 |
| Negative displacement peak (mm) | -7.66±0.64^a^ | -6.50±0.35^a^ | -9.11±1.00 ^a^ | -9.09±0.25^a^ | -8.12±3.01^a^ | -7.73±1.84^a^ |
| Positive area (N·s) | 31.31±3.07^c^ | 119.80±3.42^ab^ | 109.25±24.45^a^ | 142.93±6.22^a^ | 40.91±5.35^c^ | 56.81±3.04^bc^ |
| Negative area (N·s) | -4.31±0.47^a^ | -0.47±0.27^a^ | -1.12±1.30^a^ | -3.49±0.76^a^ | -0.03±0.02^a^ | 0.01±0.01^a^ |
| Area to positive peak (N·s) | 30.90±2.98^b^ | 57.09±1.98^ab^ | 103.50±24.19^a^ | 101.08±33.50^ab^ | 32.02±3.35^b^ | 46.50±5.97^b^ |
| Slope to positive peak (N/s) | 2.90±0.98^c^ | 12.53±0.30^a^ | 9.11±0.15^b^ | 10.77±2.15^ab^ | 3.23±0.21^c^ | 4.51±0.21^c^ |
| Force at 5 mm (N) | 6.90±0.47^c^ | 25.76±1.47^b^ | 19.45±7.67^b^ | 35.38±3.06^a^ | 5.37±0.70^c^ | 9.47±1.04^c^ |
