## Supplementary figures and images for "Solid-state fermentation of oyster mushroom by-products using *Neurospora crassa*: a sustainable approach for the development of novel meat analogues"

### Manuscript

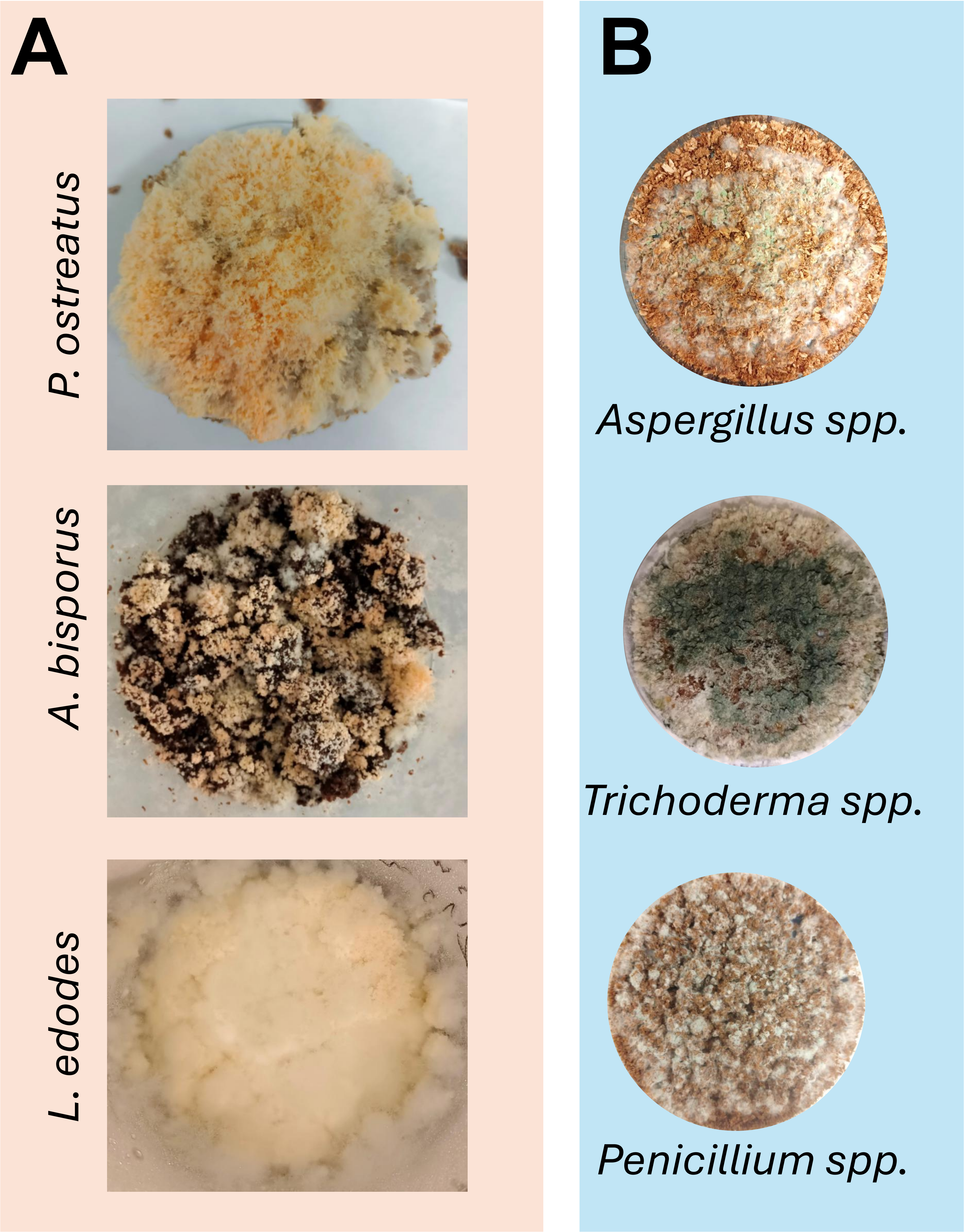
